# Five inhibitory receptors display distinct vesicular distributions in T cells

**DOI:** 10.1101/2023.07.21.550019

**Authors:** Jiahe Lu, Alisa Veler, Boris Simonetti, Timsse Raj, Po Han Chou, Stephen J. Cross, Alexander M. Phillips, Xiongtao Ruan, Lan Huynh, Andrew W. Dowsey, Dingwei Ye, Robert F. Murphy, Paul Verkade, Peter J. Cullen, Christoph Wülfing

## Abstract

T cells can express multiple inhibitory receptors. Upon induction of T cell exhaustion in response to persistent antigen, prominently in the anti-tumor immune response, many are expressed simultaneously. Key inhibitory receptors are CTLA-4, PD-1, LAG3, TIM3 and TIGIT, as investigated here. These receptors are important as central therapeutic targets in cancer immunotherapy. Inhibitory receptors are not constitutively expressed on the cell surface, but substantial fractions reside in intracellular vesicular structures. It remains unresolved to which extent the subcellular localization of different inhibitory receptors is distinct. Using quantitative imaging of subcellular distributions and plasma membrane insertion as complemented by proximity proteomics and a biochemical analysis of the association of the inhibitory receptors with trafficking adaptors, the subcellular distributions of the five inhibitory receptors were discrete. The distribution of CTLA-4 was most distinct with preferential association with lysosomal-derived vesicles and the sorting nexin 1/2/5/6 transport machinery. With a lack of evidence for the existence of specific vesicle subtypes to explain divergent inhibitory receptor distributions, we suggest that such distributions are driven by divergent trafficking through an overlapping joint set of vesicular structures. This extensive characterization of the subcellular localization of five inhibitory receptors in relation to each other lays the foundation for the molecular investigation of their trafficking and its therapeutic exploitation.

## Introduction

T cells activate in the cellular interaction with antigen presenting cells (APC). Recognition of a peptide/MHC complex on the APC surface by the T cell receptor (TCR) mediates immunological specificity and provides a strong stimulatory signal. This signal is tuned by the engagement of multiple costimulatory and coinhibitory receptors engaging ligands on the APC. An important challenge is to determine unique and overlapping functions among these coregulatory receptors. T cells express five key inhibitory receptors, CTLA-4, PD-1, LAG3, TIM3 and TIGIT [1–3]. While these receptors are also expressed in many other cell types, we focus on expression in T cells here. Inhibitory receptors can provide negative feedback in T cell activation as a default limitation of T cell activation, as is the case for CTLA-4 and PD-1 [4,5]. They can suppress T cell activation under defined physiological circumstances, for example upon persistent antigen stimulation as is the case for TIM3 and the highest levels of PD-1 expression [2]. A single T cell can express all five receptors at the same time. As the inhibitory receptors attenuate T cell activation their signaling needs to be carefully controlled. A key control element is that a substantial fraction of the cellular pool of an inhibitory receptor is kept in vesicles in the cell interior where the inhibitory receptors cannot engage their ligands on the APC surface. Regulated trafficking of vesicles containing inhibitory receptors to the cell surface in combination with endocytosis allows for control of their cell surface expression and hence function.

Vesicular localization of individual inhibitory receptors has already been characterized, that of CTLA-4 most extensively so [6,7]. Briefly, more than 80% of the cellular pool of CTLA-4 is intracellular [8]. A cytoplasmic YVKM motif allows for constitutive AP-2-mediated endocytosis when not phosphorylated, limiting CTLA-4 cell surface expression in resting T cells and allowing for an increase upon tyrosine phosphorylation during T cell activation [6,8]. Association of endocytosed CTLA-4 with LPS-responsive beige-like anchor protein (LRBA) targets CTLA-4 for lysosomal degradation [9]. Association of CTLA-4 with TRIM, LAX and Rab8 allows for regulated Golgi exit [10,11]. CTLA-4 can also be transported to the cell surface via secretory lysosomes [12] and be included in secretory microvesicles [13]. CTLA-4 trafficking is regulated by Rab5, Rab7 and Rab11 [14]. Upon T cell activation, the level of CTLA-4 at the cellular interface increases with a spatial preference for its center [15,16]. Ligand binding triggers CTLA-4 internalization to the extent that ligands can be stripped from the interacting APC [17]. A substantial part of PD-1 is located in Golgi-associated vesicles, and PD-1 accumulates at the cell surface upon T cell activation [18,19]. PD-1 can be included into ectosomes and rapidly endocytosed [20]. The majority of the cellular pool of LAG-3 is also located in vesicles overlapping with endosomal and lysosomal markers [21,22]. Upon T cell activation trafficking is diverted from lysosomes to the cell surface [22]. Substantial intracellular localization of TIM3 has been described including in Rab5 labelled vesicles and the *trans*-Golgi network, and TIM3 accumulates at the cellular interface with APCs [23,24]. However, substantial questions about inhibitory receptor localization remain unresolved. Are the subcellular distributions of different inhibitory receptors and the cellular and molecular determinants thereof distinct? Is inhibitory receptor insertion into the plasma membrane and their localization there coordinated? Here, we address these questions.

## Results

### Inhibitory receptors are enriched in the cell interior with varying degrees of clustering

To determine the subcellular distributions of inhibitory receptors, we generated fusion proteins of CTLA-4, PD-1, LAG3, TIGIT and TIM3 with GFP at their C-termini and expressed the chimeric proteins in primary T cells using retroviral transduction. Such fusion proteins have previously been validated [18,22,25–27]. To allow comparison to a large established data set, we first used CD4^+^ 5C.C7 T cell receptor (TCR) transgenic T cells [28]. We sorted transduced cells to about 600,000 GFP molecules per cell [29] and imaged them with spinning disk confocal microscopy (Figure 1A). We segmented T cells in the differential interference contrast (DIC) image, identified the midplane of a three-dimensional GFP fluorescence stack and divided the cell into their edge, comprising 30-40% or the midplane area, and their interior, the remainder (Figure 1A). For all five inhibitory receptors as compared to three receptors with established expression on the cell surface, CD2, CD28 and CD6, a significantly (p<0.01) smaller fraction of the total fluorescence was located at the cell edge as opposed to the cell interior (Figure 1B). These data suggest enhanced vesicular localization of all inhibitory receptors. By visual inspection receptor distributions displayed a different degree of clustering (Figure 1A). Using the Gini coefficient for quantification, the distribution of CTLA-4 was significantly (p<0.01) more clustered than that of any other receptors except for CD2 (Figure 1C). The LAG3 distribution, while less clustered than that of CTLA-4, was significantly (p<0.01) more clustered than that of the other three inhibitory receptors, PD-1, TIGIT and TIM3 (Figure 1C). Ripley’s K analysis of the same data as an alternate means to quantify clustering yielded the same trends (Figure S1A).

**Figure 1.**
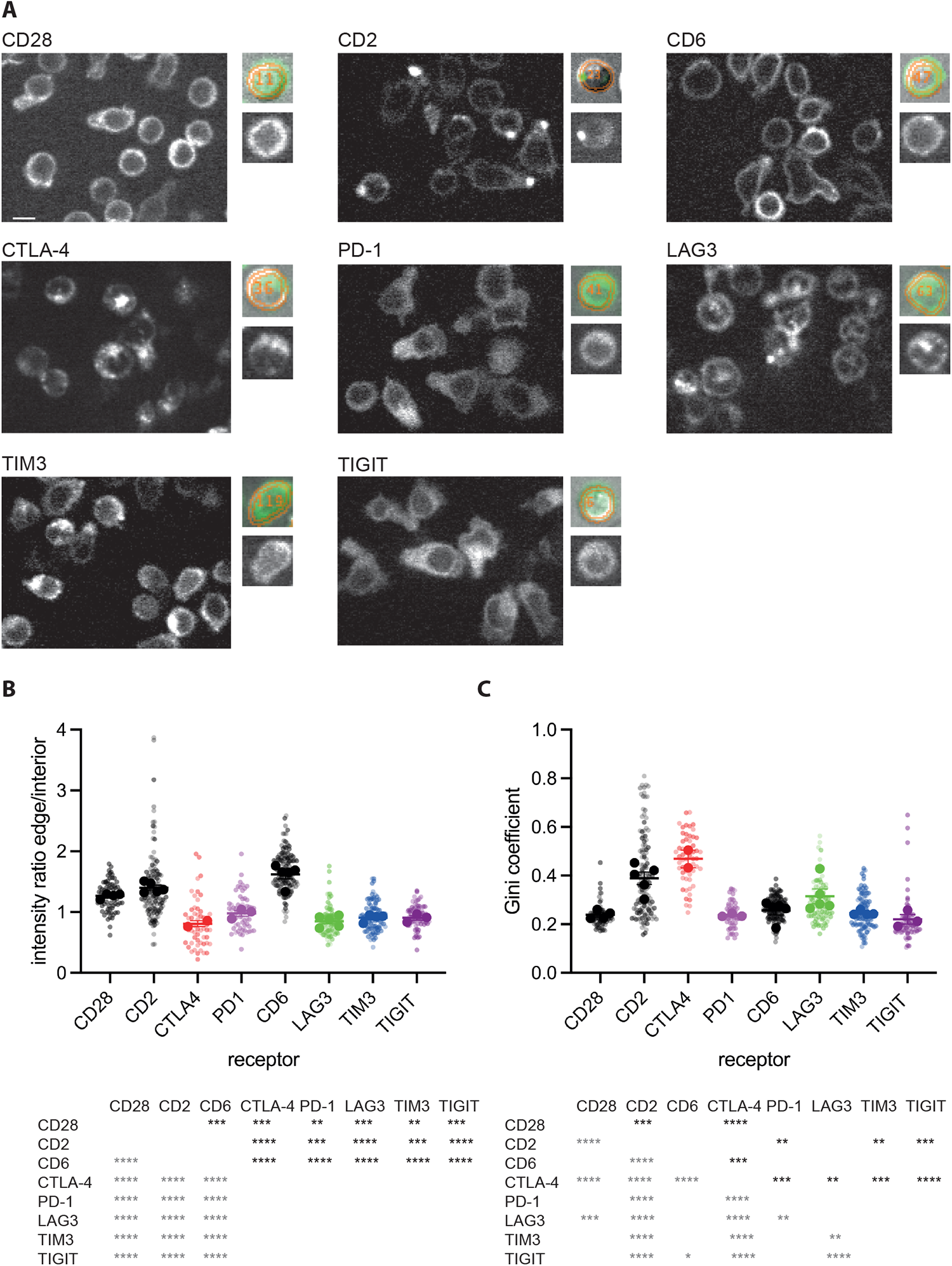
Inhibitory receptors are enriched in the interior of CD4^+^ T cells. **A** Representative spinning disk confocal midplane images of the GFP fluorescence of 5C.C7 T cells expressing a GFP fusion protein of the indicated receptor. The small images show, on the bottom, an individual cell and, on top, the matching overlay of the same GFP fluorescence in green and the analysis masks for the cell edge and interior in orange over the corresponding DIC image. Scale bar=5µm. **B** The ratio of the midplane spinning disk confocal GFP fluorescence at the T cell edge over the interior of 5C.C7 T cells expressing a GFP fusion protein of the indicated receptor as mean±SEM of run averages. Small symbols are individual cells with independent experiments denoted by color intensity. Large symbols are run averages. 56 to 141 cells per receptor from 2 to 5 independent experiments. Statistical significance of differences between receptors is given in the table below, calculated based on individual cells on top in black, based on run averages on the left in grey. **C** Gini coefficients for the same T cells as in B, given as in B. * p<0.05, ** p<0.01, *** p<0.001, **** p<0.0001; p values calculated using one-way ANOVA

Inhibitory receptors play a substantial role in the killing of tumor target cells by cytotoxic T cells (CTL). We, therefore, repeated the localization analysis in CD8^+^ CL4 TCR transgenic T cells (Figure 2A) [30]. We use the interaction of CL4 CTL with Renca renal carcinoma cells as a model to investigate tumor cell cytolysis. Similar to 5C.C7 CD4^+^ T cells, a significantly (p<0.05) smaller fraction of the total fluorescence of all five inhibitory receptors as compared to that of the TCR was located at the cell edge as opposed to the cell interior (Figure 2B). Using the Gini coefficient, the distribution of CTLA-4 was again more significantly (p<0.0001) clustered than that of the other four inhibitory receptors and that of the TCR (Figure 2C). The distribution of LAG3, while less clustered than that of CTLA-4, was significantly more clustered (p<0.05) than that of the TCR, PD-1, TIGIT and TIM3 (Figure 2B). The same trends were seen using Ripley’s k analysis (Figure S2A).

**Figure 2.**
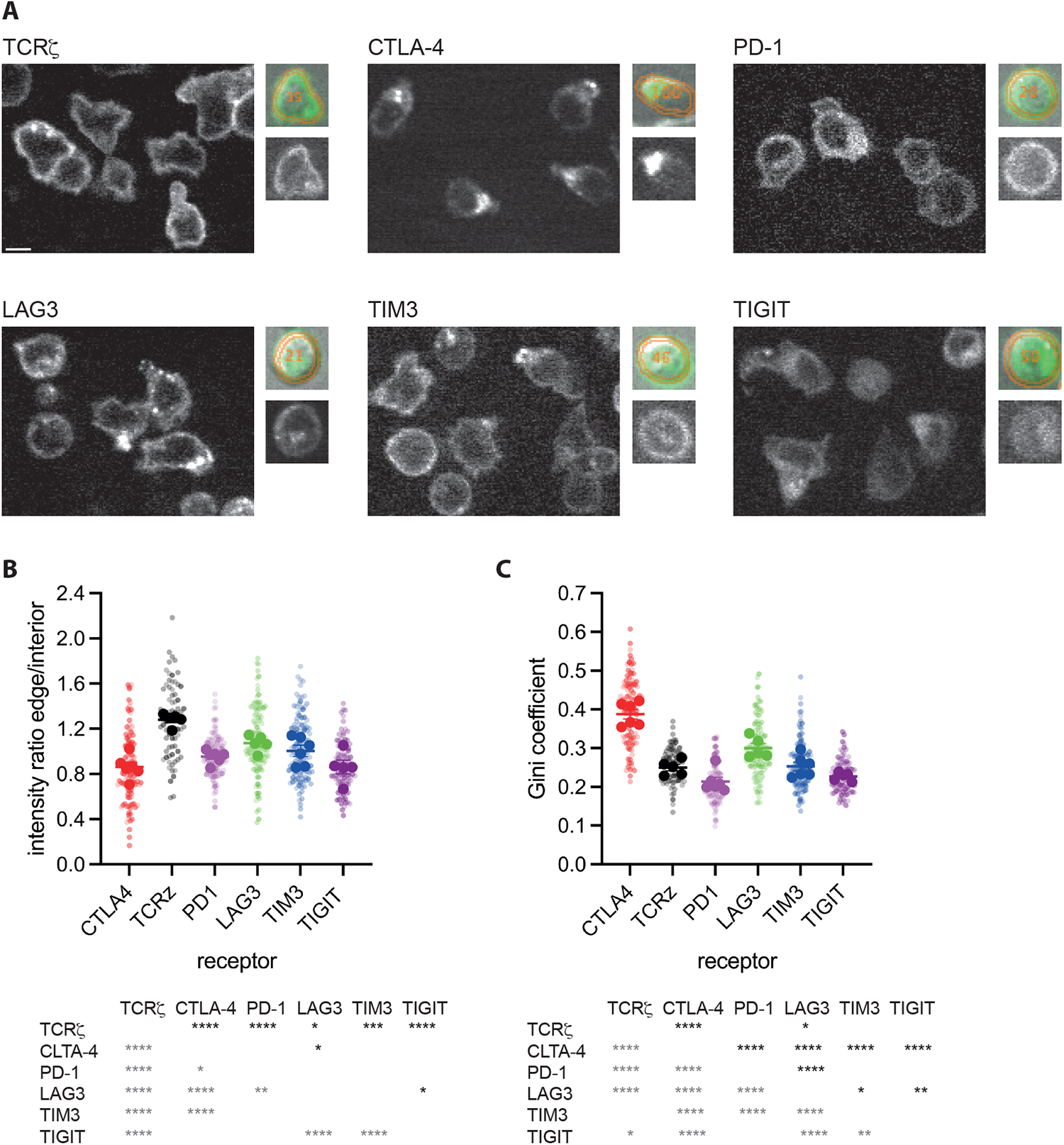
Inhibitory receptors are enriched in the interior of CD8^+^ T cells. **A** Representative spinning disk confocal midplane images of the GFP fluorescence of CL4 T cells expressing a GFP fusion protein of the indicated receptor. The small images show, on the bottom, an individual cell and, on top, the matching overlay of the same GFP fluorescence in green and the analysis masks for the cell edge and interior in orange over the corresponding DIC image. Scale bar=5µm. **B** The ratio of the midplane spinning disk confocal GFP fluorescence at the T cell edge over the interior of CL4 T cells expressing a GFP fusion protein of the indicated receptor as mean±SEM of run averages. Small symbols are individual cells with independent experiments denoted by color intensity. Large symbols are run averages. 77 to 131 cells per receptor from 5 to 6 independent experiments. Statistical significance of differences between receptors is given in the table below, calculated based on individual cells on top in black, based on run averages on the left in grey. **C** Gini coefficients for the same T cells as in B, given as in B.

To provide an estimate of which fraction of the inhibitory receptors is located at the plasma membrane versus the cell interior, we acquired a smaller image data set of live CL4 CTL expressing TCRζ or the five inhibitory receptors right at the resolution limit of confocal microscopy (Figure 3A). Even though the plasma membrane thickness is smaller than this resolution limit, these images allowed us to generate an automated image analysis routine approximating the fluorescence signal associated with the plasma membrane. Using this routine, only 7-28% of the cellular pool of the five inhibitory receptors was associated with the plasma membrane, significantly (p<0.01) less than the 59±2% of the TCR, a stimulatory receptor that nevertheless also continuously cycles through vesicular compartments [31,32](Figure 3B). As an alternate approach, we activated T cells with beads coated with antibodies against CD3 and CD28 for 48h and stained them at the cell surface and after permeabilization in the entire cell for the endogenous amounts of the five inhibitory receptors and CD45 as a plasma membrane marker (Figure 3C). While cell surface expression accounted for the majority of all CD45, only 9-26% of the cellular pools of the inhibitory receptors CTLA-4, TIM3 and TIGIT was expressed at the cell surface (Figure 3D), in fairly close agreement with the image analysis data. In contrast, all of PD-1 and a majority of LAG-3 were expressed at the cell surface. As PD-1 is moved to the cell surface upon lymphocyte activation [19,33], continuous TCR engagement in generating the activated T cells used for staining may have triggered PD-1 translocation to the plasma membrane here. In support, PD-1 is largely intracellular in the same T cells in the absence of TCR ligation (Figure S3). Acute TCR stimulation similarly triggers translocation of LAG3 to the cell surface [22].

**Figure 3.**
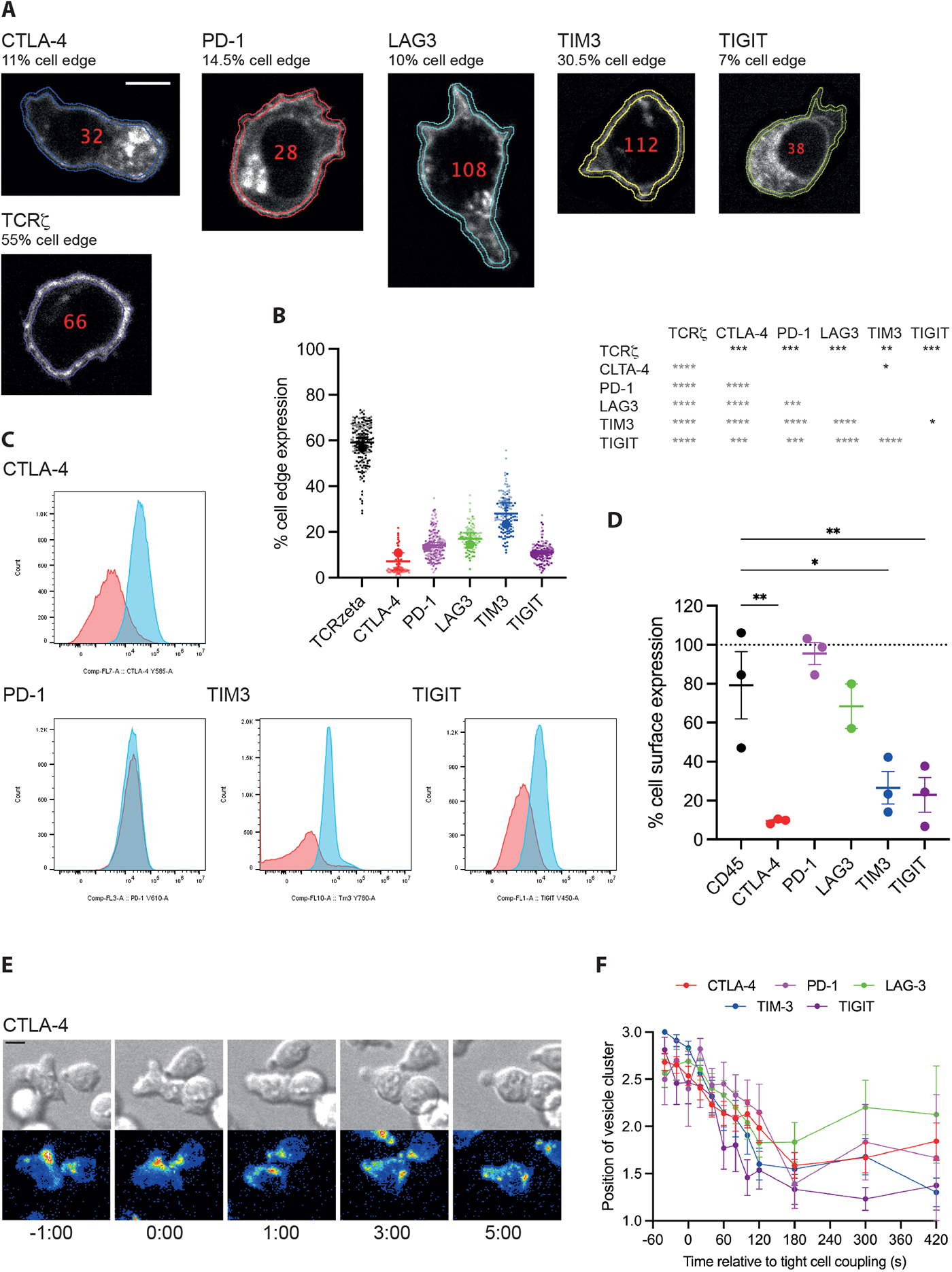
Inhibitory receptors are enriched in the interior of CD8^+^ T cells. **A** Representative spinning disk confocal midplane images of the GFP fluorescence of CL4 T cells expressing a GFP fusion protein of the indicated receptor. Scale bar=5µm. **B** The fraction of GFP fluorescence associated with the cell edge in CL4 T cells expressing a GFP fusion protein of the indicated receptor. Large symbols are averages of independent experiments with mean±SEM and color intensity denoting matched experiments. Small symbols are the constituent single cell measurements with independent experiments denoted by color intensity. 77 to 131 cells per receptor from 2 independent experiments. Statistical significance of differences between receptors is given in the table to the right, calculated based on run averages on top in black, based on individual cells on the left in grey. **C** Representative flow cytometry data of CD8^+^ T cells stimulated for 48h stained for the indicated receptor on the cell surface (red) or in the entire cell (blue). **D** Fraction of endogenous inhibitory receptor expressed on the cell surface based on flow cytometry analysis (C) given as averages of independent experiments with mean±SEM. 3 to 6 replicates from 3 independent experiments. **E** Representative time lapse image sequence of a CL4 CTL expressing CTLA-4-GFP activating in the interaction with a Renca target cell incubated with 2µg/ml HA peptide. DIC images on top, maximum projection images of the three-dimensional GFP fluorescence in a rainbow false color scale on the bottom, time given in minutes. Scale bar=5µm. **F** Average position of a single cluster of vesicles in cell couples of CL4 CTL expressing a GFP-fusion protein of the indicated receptor with a Renca target cell incubated with 2µg/ml HA peptide over time as mean±SEM. 1 is at the cellular interface, 3 is at the distal pole. 15 to 41 cell couples per receptor from 3 to 4 independent experiments. * p<0.05, ** p<0.01, *** p<0.001, **** p<0.0001; p values calculated using one-way ANOVA

Upon binding of a T cell to an antigen presenting cell (APC) the MTOC reorients within two minutes from behind the nucleus to the cellular interface [34,35]. To determine whether the distributions of the inhibitory receptors, as expected from vesicular localization, display a similar relocalization, we activated CL4 CTL with Renca tumor target cells incubated with a high concentration, 2µg/ml, of the Influenza hemagglutinin peptide 518-526 and imaged the interaction with spinning disk confocal microscopy. A reorientation of vesicular clusters, to the extent that they could be detected, from behind the nucleus toward the cellular interface was consistently observed (Figure 3E, F). Together, these data are consistent with the suggestion that a substantial fraction of the cellular pool of all five inhibitory receptors resides in vesicles with distributions that combine some shared features with pronounced differences between different inhibitory receptors.

### Vesicles transporting inhibitory receptors are inserted into the T cell plasma membrane upon T cell activation with distinct frequency and kinetics resulting in distinct interface distributions

As inhibitory receptors are localized in vesicles, one should be able to detect the insertion of such vesicles into the T cell plasma membrane upon T cell activation. To test this suggestion, we imaged the activation of CL4 T cells on glass cover slips coated with the stimulatory anti-TCR antibody 145-2C11 with or without the extracellular domain of the costimulatory ligand ICAM-1 using total internal reflection (TIRF) microscopy. CL4 T cells spread on the cover slips as an indication of activation (Figure 4A) and events consistent with vesicle insertion into the T cell plasma membrane, rapid appearance of fluorescence followed by lateral dispersion, were readily detectable (Figure 4B). We used a machine learning-based analysis routine to automatically detect such events during CL4 T cell activation. To capture dynamics, we tabulated insertion events in time windows of 100 frames, corresponding to about 30s. The overall number of insertion events differed between inhibitory receptors, being the smallest for CTLA-4 and the highest for LAG3 (Figure 4C). Dynamics also differed. While the number of insertion events for CTLA-4 consistently increased over time, it consistently decreased for TIGIT (Figure 4D). For PD-1, LAG3 and TIM3 an initial burst of insertion events within the first 100 frames was followed by largely stable insertion thereafter (Figure 4D).

**Figure 4.**
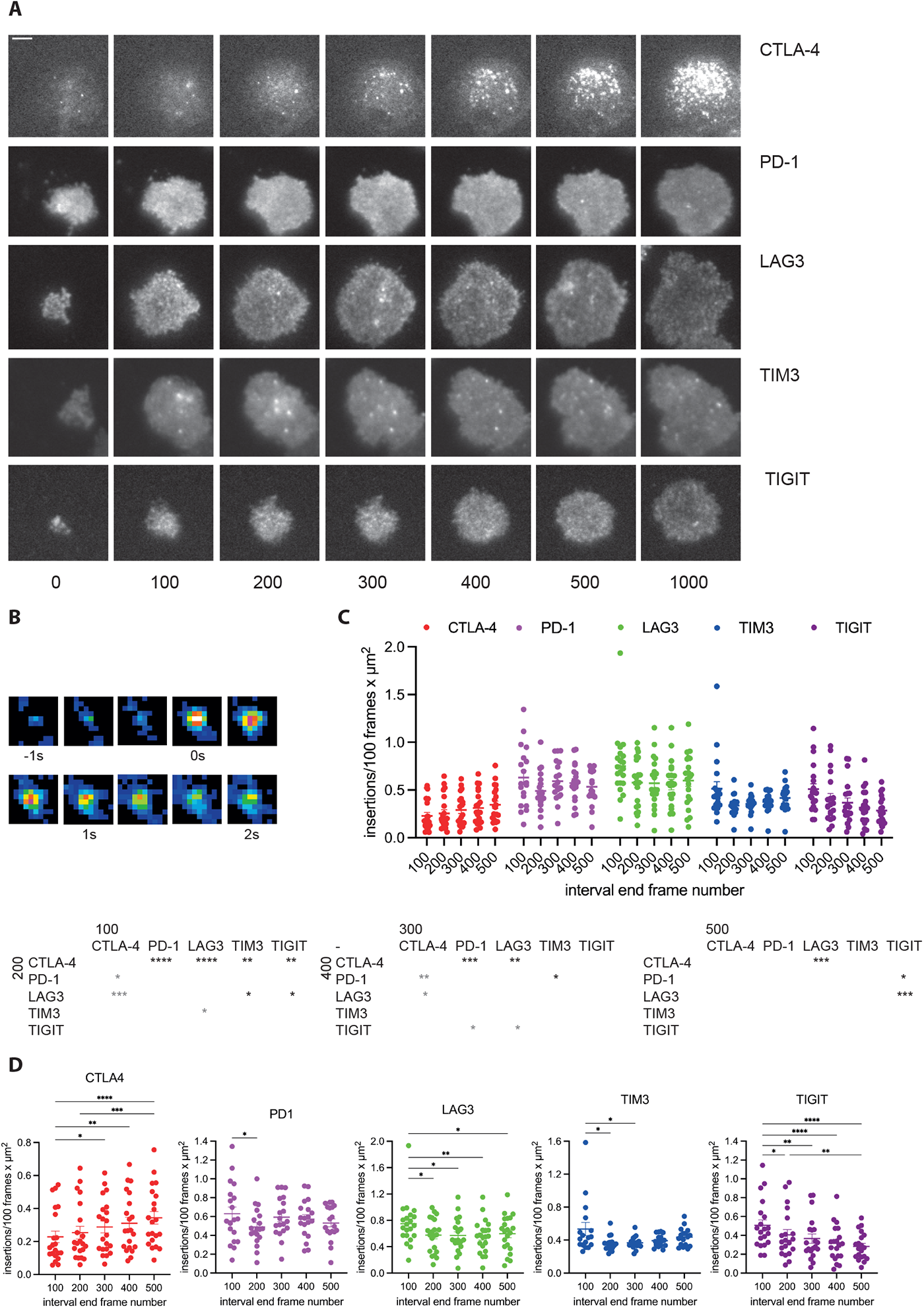
Inhibitory receptors are inserted into the plasma membrane during T cell activation. **A** Representative time lapse TIRF images of the GFP fluorescence of CL4 CTL expressing a GFP fusion protein of the indicated receptor activated on glass cover slips coated with anti-CD3ε with or without ICAM-1. Time given in seconds. Scale bar=2µm **B** Time lapse TIRF image sequence of the GFP fluorescence in a false color rainbow scale of a single membrane insertion event at frame rate. **C** For the experiments shown in A, number of insertion events per 100 frames and µm^2^ for the indicated receptors as individual cells plus mean±SEM. Time is denoted by the number of the last frame in the time interval. 100 frames correspond to 30s. 19 to 21 cells from 4 to 6 independent experiments. Statistical significance of differences between receptors is given in the tables below for each of the five time windows as indicated. **D** The same data as in C displayed in separate panels for the individual receptors with statistical differences between time windows. * p<0.05, ** p<0.01, *** p<0.001, **** p<0.0001; p values calculated using one-way (D) and two-way (C) ANOVA

To determine whether the divergent insertion of vesicles carrying different inhibitory receptors resulted in distinct inhibitory receptor distributions across the T cell coverslip interface, we performed a line scan analysis of the TRIF data where we measured the fluorescence intensity in ten equally sized segments of a line of a width of 4.5µm across the entire interface. The CTLA-4 distribution was focused consistently more to the center of the interface (Figure 5A-D). While LAG3 was slightly focused on the interface center at an early time point (first 100 frames of T cell spreading), it was uniquely enriched at the interface edge at the late time point (≥ 5min after initial spreading)(Figure 5A-D). Distributions of PD-1, TIGIT and TIM3 were similar with a slight focus on the interface center that decreased over time (Figure 5A-D). To confirm that the divergent distributions derived from the single-color analysis of the five inhibitory receptors could be recapitulated in a two-color experiment, we imaged CL4 CTL expressing a fusion protein of LAG3 with tdTomato with either CTLA-4-GFP or TIM3-GFP (Figure 5E). Consistent with the selective focus of the CTLA-4 distribution on the interface center, colocalization of LAG3 with CTLA-4 was less than that with TIM3 (Figure 5F). Consistent with the selective translocation of LAG-3 to the interface edge, colocalization between LAG3 and TIM3 declined over time (Figure 5F). The two selected two-color experiments thus validate the single-color analyses.

**Figure 5.**
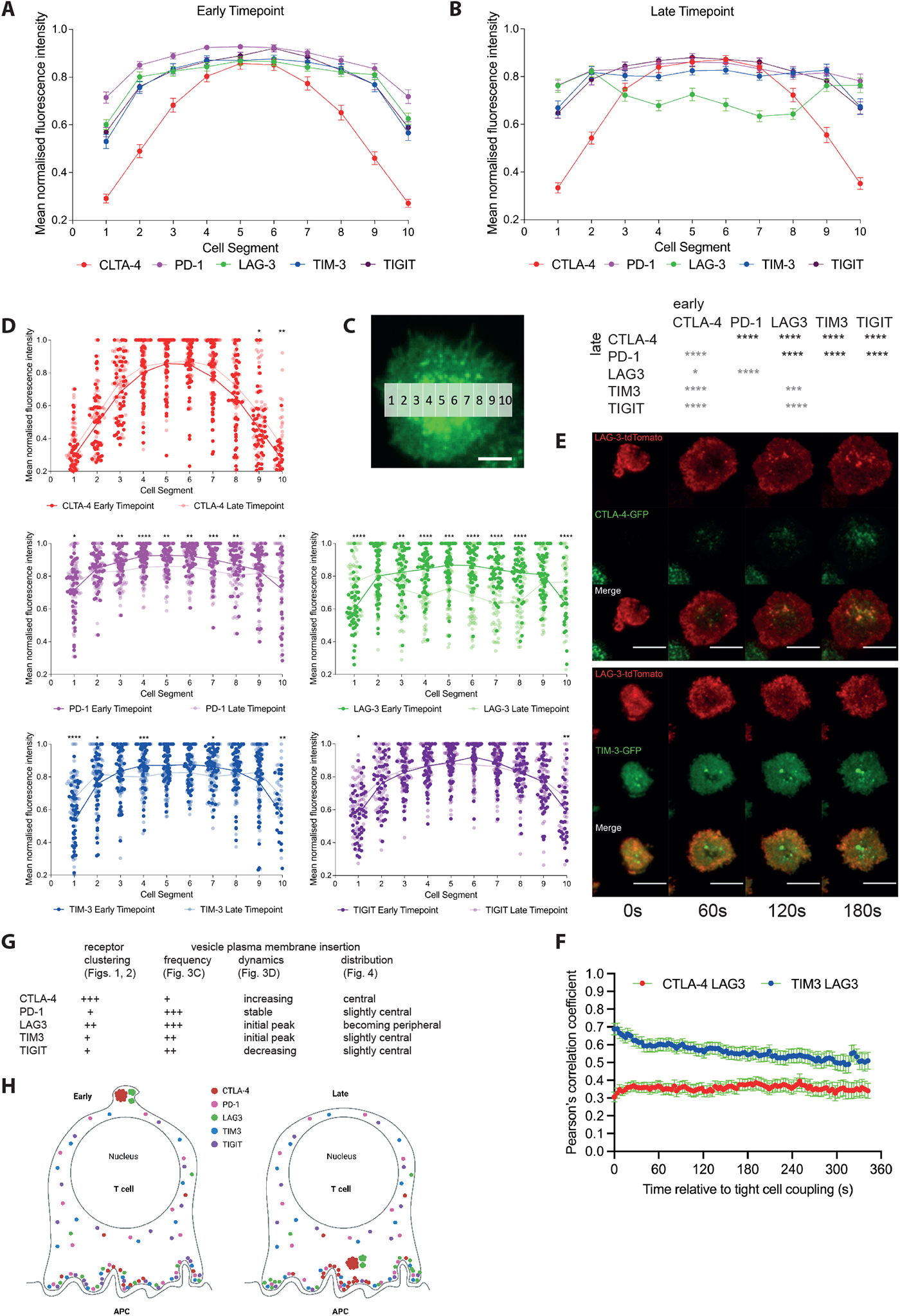
Inhibitory receptors are distinctly distributed in the plasma membrane during T cell activation. **A-C** CL4 CTL expressing a GFP fusion protein of the indicated receptor were activated on glass cover slips coated with anti-CD3ε and ICAM-1 and imaged by TIRF. Given is the GFP fluorescence intensity in ten equally sized segments of a line scan with a line width of 5µm across the entire interface as mean±SEM normalized as 1 for the segment with the highest intensity within 30s (A) and >5min (B) of CTL spreading. 50 CTL analyzed per receptor from 4 to 6 independent experiments. Statistical significance of differences between receptors is given in the table separately for the early (in grey on the left) and late (in black on top) time points. C The representative image shows the 10 measurement regions. Scale bar=5µm. **D** The same data as in A are displayed separately for each inhibitory receptor with statistical significance of differences between early and late time point in each line scan segment. **E** Representative time lapse TIRF images of the GFP and tdTomato fluorescence of CL4 CTL expressing the indicated fluorescent protein fusion protein of the indicated receptors activated on glass cover slips coated with anti-CD3ε and ICAM-1. Time given in seconds. Scale bar=10µm. **F** For the experiments shown in E and the indicated combinations of inhibitory receptors, Pearson’s correlation coefficients for GFP and tdTomato fluorescence across the entire interface as mean±SEM over time. **G, H** Summary of imaging data from Figs. 1 to 5, as a table (G) and graphically as created with Biorender.com (H). * p<0.05, ** p<0.01, *** p<0.001, **** p<0.0001; p values calculated using two-way ANOVA (A) and Wilcoxon matched-pairs rank test (B).

In summary, across all imaging assays the five inhibitory receptors displayed distributions consistent with a predominant vesicular localization. This contrasted with the stronger enrichment of the TCR, the costimulatory receptors CD28 and CD2, and CD6 at the cell edge. When analyzed in conjunction cross all imaging assays, distinct localization profiles emerged for all inhibitory receptors (Figure 5G, H). CTLA-4 stood out with the highest degree of clustering, the lowest rate of vesicular plasma membrane insertion, the only insertion rate increasing over time and strongly central interface localization. PD-1 in contrast combined little clustering with a high, stable rate of vesicular insertion. LAG3 clustering was second highest to that of CTLA-4 and high-rate vesicular insertion led to a unique accumulation at the interface periphery. The distributions of TIGIT and TIM3 were most similar with little clustering and intermediate rate vesicular insertion with a slight preference for the interface center. Yet, while the rate of vesicular insertion of TIGIT was steadily declining, that of TIM3 was slightly increasing after a rapid decline of a pronounced initial peak.

### The proteomes of vesicles harboring CTLA-4, LAG3 and TIM3 are distinct

To corroborate distinct subcellular localization of inhibitory receptors in molecular terms, we used APEX2 proteomics [36,37], a biotin-based proximity-labelling approach, to label the lumen of vesicles containing inhibitory receptors. We tagged all five inhibitory receptors with the engineered peroxidase APEX2 at the extracellular domain, the domain pointing into the vesicle lumen, and with GFP at the cytoplasmic domain. These proteins were retrovirally expressed in primary CL4 CTL and the transduced CTL sorted for expression of the tagged receptors. Only expression of the APEX2-tagged versions of CTLA-4, LAG3 and TIM3 were sufficiently high to allow determination of the luminal vesicular proteome. Using streptavidin labeled with Alexa Fluor 568 to detect proteins biotinylated by APEX2, we confirmed that APEX2 labelling was specific for structures enriched in the inhibitory receptors (Figure S4). Using an unbiased analysis method based directly on peptide counts in mass spectrometry, the vesicular proteomes of CTLA-4, LAG3 and TIM3 were reproducible and distinct (Figure 6A). Selective enrichment of CTLA-4 and LAG3 in the respective proteomes supports specificity of the analysis (Figure 6B). Enrichment of the transferrin receptor (Tfrc), CD45 (Ptprc) and Transmembrane p24 trafficking protein 10 (Tmed10) in all three vesicular proteomes supports endocytic trafficking of all three receptors as an overlapping part of their vesicular localization (Figure 6B). Thus, these proteomics data corroborate the imaging data showing that inhibitory receptors reside in vesicular populations that differ between the receptors. The shared enrichment of some vesicular proteins suggests that the receptors traffic at least partially through the same vesicle types, possibly with a divergent degree of enrichment. The identification of proteins uniquely enriched for individual receptors allows for the possibility of unique vesicle types harboring these receptors. However, the detected luminal proteins didn’t further allow the identification of specific types of vesicles harboring the inhibitory receptors, likely because the transport machineries mediating vesicular identify are largely bound to the cytoplasmic, not the luminal side of the vesicle membranes.

**Figure 6.**
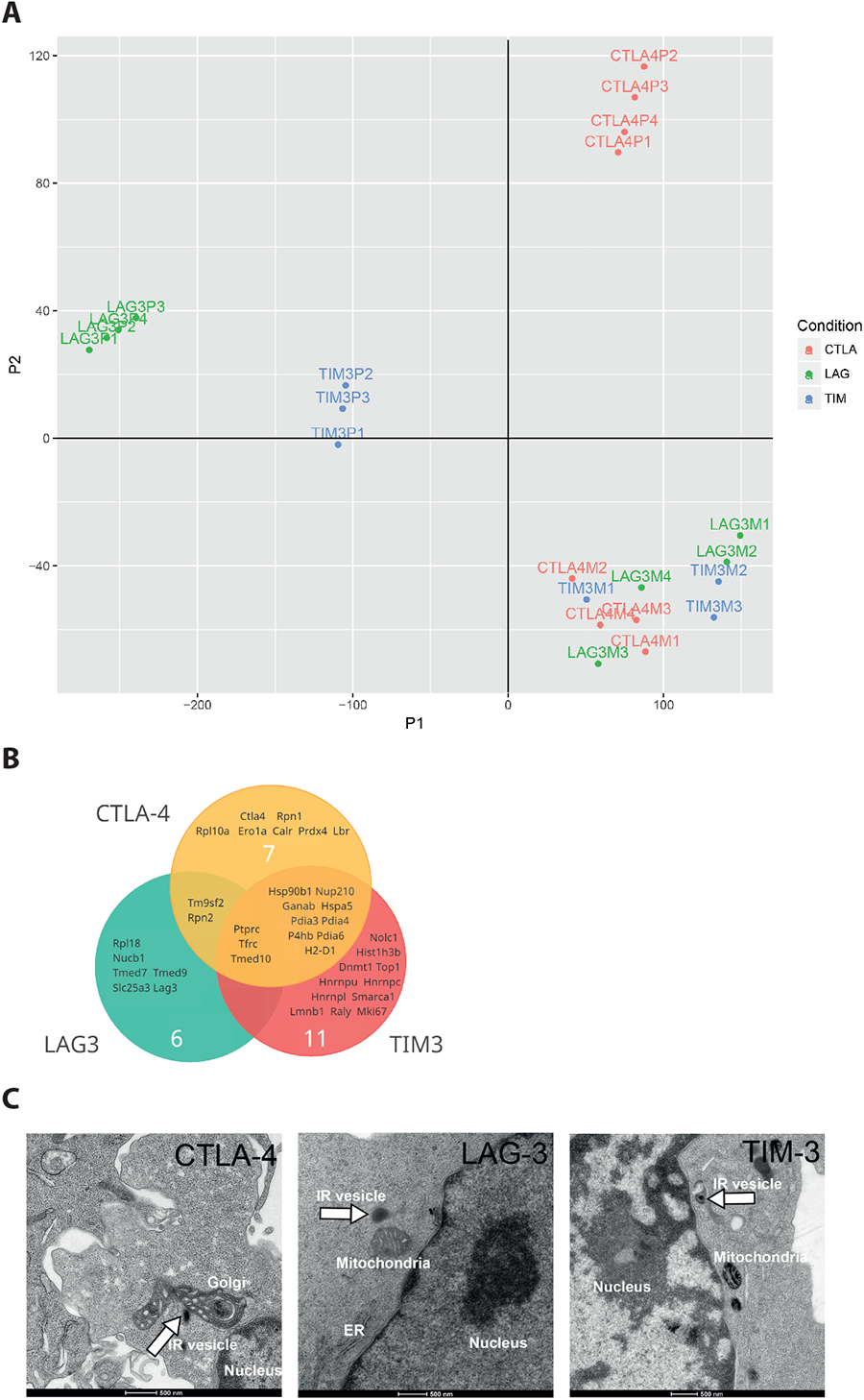
The vesicular proteomes of CTLA-4, LAG3 and TIM3 are distinct. **A** t-SNE plot of raw mass spectrometry peptide data after APEX2-mediated proximity biotinylation of proteins in the lumen of vesicles harboring APEX2 fusion proteins of CTLA-4, LAG3 and TIM3 expressed in CL4 CTL. Each dot is an independent experimental repeat labelled as P1 to P4 for the APEX2 experiments and as M1 to M4 for the no-APEX2 control for endogenously biotinylated proteins. **B** Gene names of proteins detected with a global false discovery rate of p<0.05 in the lumen of vesicles expressing APEX2 fusions of CTLA-4, LAG3 and TIM3 and combinations thereof. **C** Examples of electron micrographs of CL4 CTL where vesicles expressing APEX2 fusion proteins of CTLA-4, LAG3 or TIM3 as indicated are labelled with an APEX2-mediated electron dense precipitate as marked with a white arrow. Scale bar=0.5µm.

Our extracellular fusion of inhibitory receptors to APEX2 allows for the labelling of vesicles harboring the receptors in electron micrographs [37]. We could detect vesicles labelled with electron dense material (Figure 6C). These data are consistent with vesicular localization of inhibitory receptors. However, in a blind analysis of the entire data, specificity of electron-dense labelling was not sufficiently high to allow quantification and detailed characterization of vesicle types by their morphology.

### CTLA-4 associates more extensively with lysosomes

To further characterize vesicles harboring inhibitory receptors, we incubated CL4 CTL expressing GFP fusion proteins of the inhibitory receptors for 10min with Renca target cells incubated with a high concentration, 2 µg/ml, of HA agonist peptide. We then stained the cell couples with an antibody against EEA1 to label early endosomes, and with Lysotracker and Mitotracker to label lysosomes and mitochondria, respectively. We automatically segmented the cytoplasm of individual T cells (Figure S5A) and determined the Pearson’s correlation coefficient between inhibitory receptor and vesicular marker distributions (Figure 7). TIGIT-GFP expression was not high enough to allow an effective execution of such analysis. Correlation between the inhibitory receptors and individual vesicular marker was moderately positive, consistent with a distribution of inhibitory receptors across multiple vesicle types. Association of CTLA-4 with lysosomes was significantly (p<0.05) higher than that of the other three receptors investigated (Figure 7A, B). As secretory granules of CTL are lysosomally derived, these data suggest that CTLA-4 selectively traffics with secretory granules. Association of inhibitory receptors with endosomes was overall highest for the vesicular markers (Figure 7C, D), suggesting consistent internalization of inhibitory receptors from the plasma membrane. In an exploratory experiment, we also investigated endosomal association at a single 20-minute time point of T cell activation (Figure S5B, C). At this time CTLA-4 was significantly (p<0.05) more strongly associated with endosomes than LAG3 and TIM3 reaching an average Pearson’s correlation coefficient of 0.45±0.1. While a further investigation of the time dependence of vesicular associations is beyond the scope of this manuscript, the data suggest that vesicular associations vary over time and that such variation differs between the inhibitory receptors. We also investigated association of inhibitory receptors with mitochondria, largely as a negative control but also as the APEX proteomics found enrichment of mitochondrial proteins such as the phosphate carrier Slc25a3 in the lumen of vesicles carrying LAG3. Colocalization with mitochondria was minimal as expected (Figure 7E, F). Overall, the colocalization data identified a preferential association of CTLA-4 with lysosomally derived vesicles with potentially enhanced endocytosis late in T cell activation. Substantial differences in the localization of PD-1, LAG3 and TIM3 were not detected, possibly because these receptors are distributed across multiple vesicle types.

**Figure 7.**
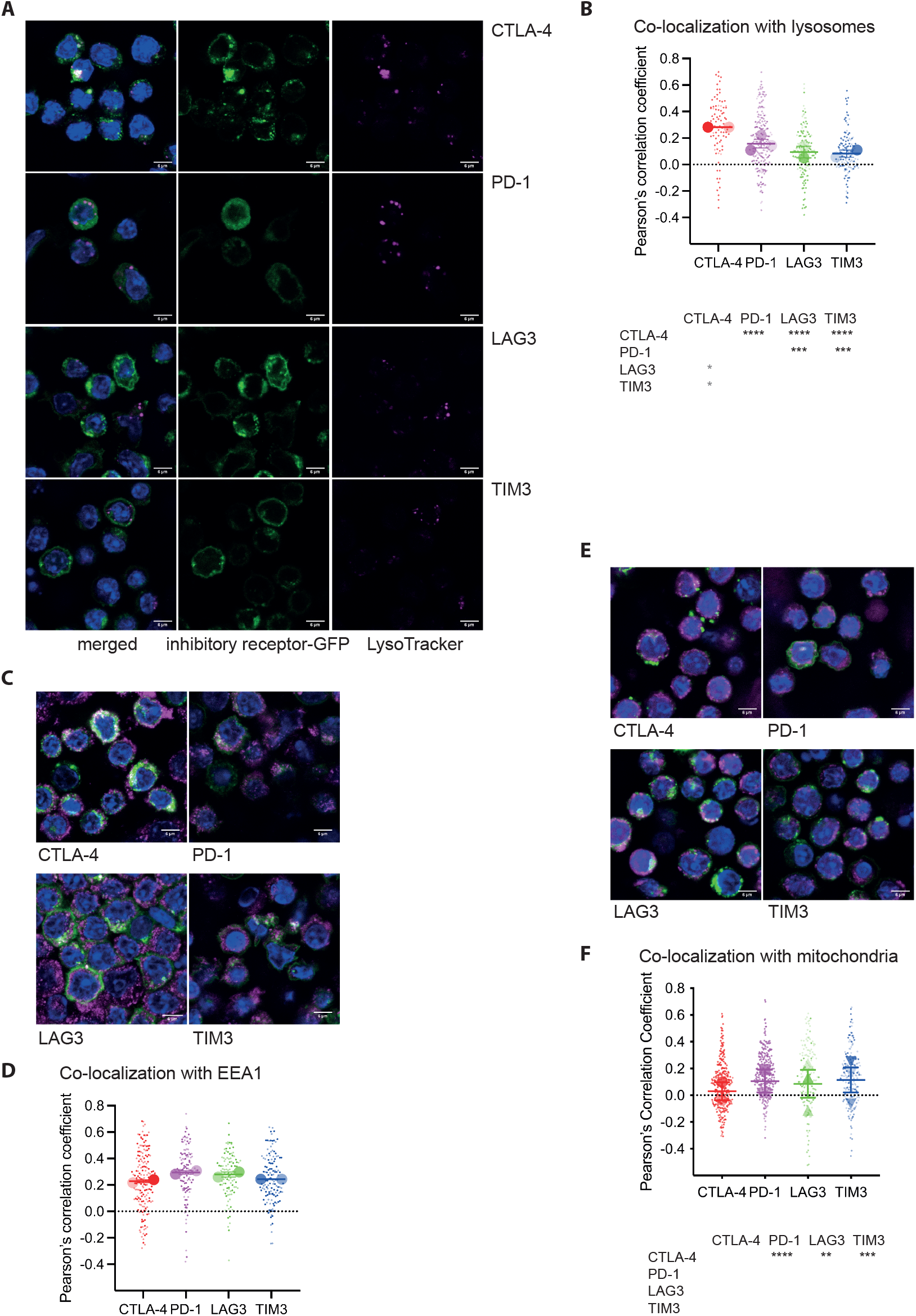
CTLA-4 is selectively enriched in lysosomally derived vesicles. **A** CL4 CTL expressing GFP fusion proteins of the indicated inhibitory receptors and incubated with LysoTracker Red DND-99 were activated by incubation with Renca target cells in the presence of 2µg/ml HA agonist peptide for 10min, fixed and stained for nuclei with Hoechst 33258. Given are confocal z-stack midplane images of the LysoTracker fluorescence (magenta), the GFP fluorescence (green) and merged images of all fluorescence data (Hoechst 33258 in blue). Scale bar=6µm. **B** For the experiment in A, Pearson’s correlation coefficients of the GFP and LysoTracker fluorescence distributions in the cytoplasm of individual CL4 CTL segmented from three-dimensional fluorescence data. Small symbols are individual cells with independent experiments denoted by color intensity. Large symbols are run averages. 91 to 207 cells per receptor from 2 to 3 independent experiments. Statistical significance of differences between receptors is given in the table below, calculated based on individual cells on top in black, based on run averages on the left in grey. **C** CL4 CTL expressing GFP fusion proteins of the indicated inhibitory receptors were activated by incubation with Renca target cells in the presence of 2µg/ml HA agonist peptide for 10min, fixed and stained with anti-EAA1 and for nuclei with Hoechst 33258. Given are merged confocal z-stack midplane images of the anti-EAA1 (magenta), GFP (green) and Hoechst 33285 (blue) fluorescence. Scale bar=6µm. **D** For the experiment in C, Pearson’s correlation coefficients of the GFP and EAA1 fluorescence distributions in the cytoplasm of individual CL4 CTL segmented from three-dimensional fluorescence data. Small symbols are individual cells with independent experiments denoted by color intensity. Large symbols are run averages. 101 to 166 cells per receptor from 2 independent experiments. There were no significant differences between the inhibitory receptors. **E** CL4 CTL expressing GFP fusion proteins of the indicated inhibitory receptors and incubated with MitoTracker Red CMXRos were activated by incubation with Renca target cells in the presence of 2µg/ml HA agonist peptide for 10min, fixed and stained for nuclei with Hoechst 33258. Given are merged confocal z-stack midplane images of the MitoTracker (magenta), GFP (green) and Hoechst 33285 (blue) fluorescence. Scale bar=6µm. **F** For the experiment in E, Pearson’s correlation coefficients of the GFP and MitoTracker fluorescence distributions in the cytoplasm of individual CL4 CTL segmented from three-dimensional fluorescence data. Small symbols are individual cells with independent experiments denoted by color intensity. Large symbols are run averages. 151 to 298 cells per receptor from 2 to 3 independent experiments. Statistical significance of differences between receptors is given in the table below, calculated based on individual cells on top in black, based on run averages on the left with no significant differences.

### Association with regulators of receptor transport is largely shared across the five inhibitory receptors

The cytoplasmic domains of the inhibitory receptors contain multiple sorting motifs conserved between mouse and human that may mediate the selective binding of key regulators of receptor trafficking (Figure 8A). To determine the ability of the cytoplasmic domains of the inhibitory receptors to recruit regulators of receptor trafficking, we expressed GFP fused to the cytoplasmic domains of the inhibitory receptors in HEK-293T cells, precipitated the GFP fusion proteins and probed for associated regulators of receptor trafficking. A punctuate distribution of the inhibitory receptor cytoplasmic domain-GFP proteins suggested some vesicular association (Figure 8B). However, we could not selectively precipitate a number of key regulators of receptor trafficking with these fusion proteins (Figure 8C-E).

**Figure 8.**
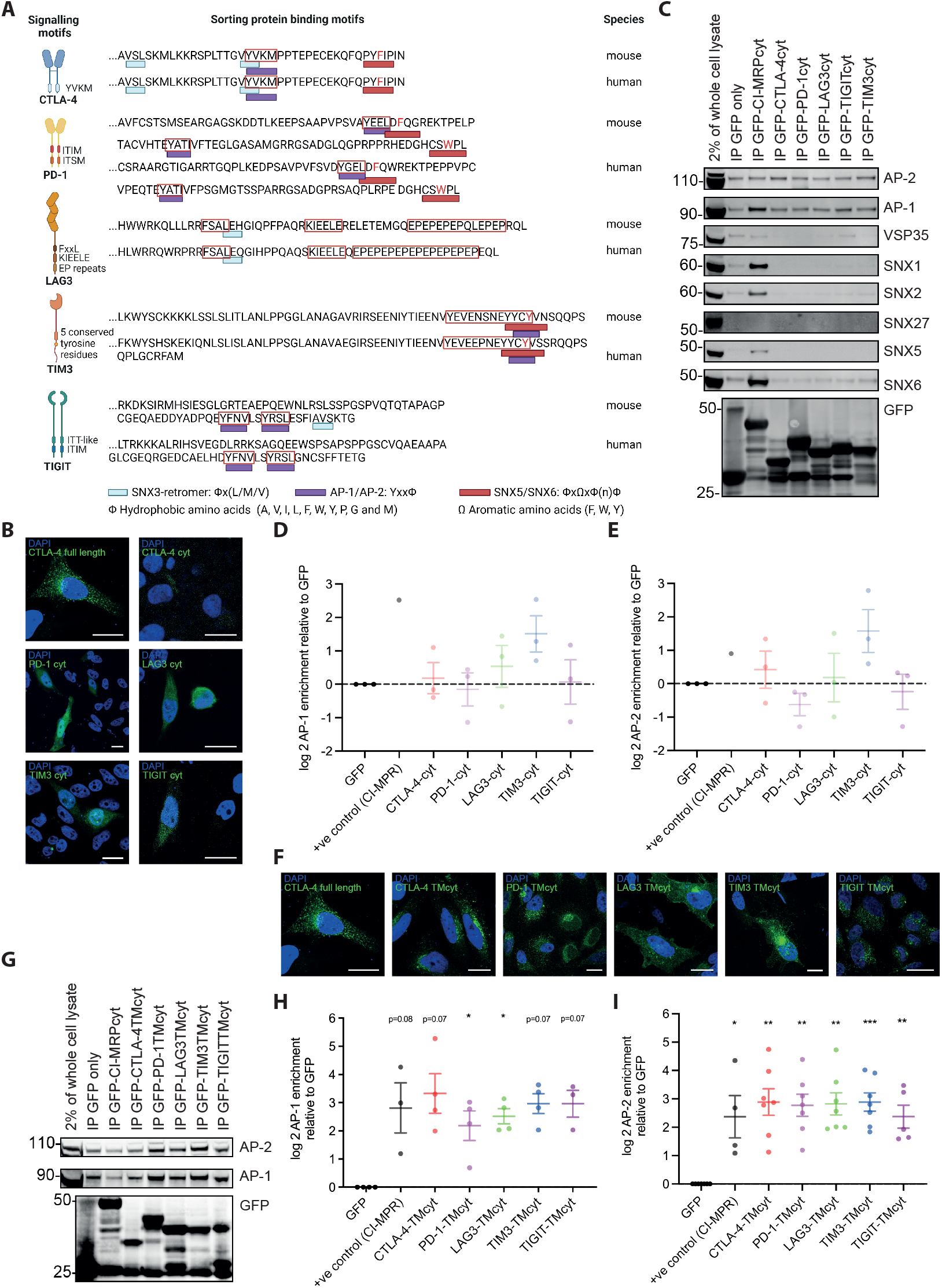
The cytoplasmic and transmembrane domains of the inhibitory receptors associate with AP-1 and AP-2. **A** Protein sorting motifs in the cytoplasmic domains of the inhibitory receptors. Figure panel created with BioRender.com. **B** Representative images of HEK293T cells expressing fusion proteins of the cytoplasmic domains of the inhibitory receptors and of full length CTLA-4 for comparison as indicated. Scale bar=10µm. **C** Representative Western Blots of GFP pull down experiments from the HEK293T cells in B as indicated on top with antibodies against vesicular markers and GFP as a control for fusion protein expression as indicated on the right. Full size blots in Figure S6. **D, E** Log2 of enrichment of AP-1 (D) and AP-2 (E) at the cytoplasmic domains of the indicated receptors fused to GFP relative to GFP only. 3 independent experiments. **F** Representative images of HEK293T cells expressing fusion proteins of the cytoplasmic and transmembrane domains of the inhibitory receptors and of full length CTLA-4 for comparison as indicated. Scale bar=10µm. **G** Representative Western Blot of GFP pull down experiments from the HEK293T cells in F as indicated on top with antibodies against AP-1, AP-2 and GFP as a control for fusion protein expression as indicated on the right. Full size blots in Figure S6. **H, I** Log2 of enrichment of AP-1 (H) and AP-2 (I) at the cytoplasmic and transmembrane domains of the indicated receptors fused to GFP relative to GFP only. 3 to 7 independent experiments. * p<0.05, ** p<0.01, *** p<0.001 relative to GFP only; p values calculated using one-way ANOVA

To strengthen association with vesicular structures, we generated fusion proteins of GFP followed by the transmembrane and cytoplasmic domains of the inhibitory receptors and repeated the pull-down experiments. The distribution of these fusion proteins appeared more punctuate (Figure 8F), consistent with enhanced vesicular association. With these chimeras we detected the association of the inhibitory receptors with AP-1, AP-2 (Figure 8H-I), the ESCPE-1 subunits SNX1/2/5/6 [38] (Figure 9A-E) and the core Retromer subunit VPS35 [39] (Figure 9F, G). The transmembrane/cytoplasmic domain constructs of all five receptors efficiently associated with AP-1 and AP-2 (Figure 8H-I), suggesting that all receptors traffic through key conserved secretory and endocytic routes. Association with all sorting nexins and VPS35 was found for all inhibitory receptors, yet it was consistently highest for CTLA-4 without the differences between the inhibitory receptors reaching statistical significance (Figure 9). CTLA-4 association reached levels close to those of the cytoplasmic domain of the cation-independent mannose-6-phosphate receptors as a positive control for ESCPE-1-dependent retrograde transport. These data suggest that all inhibitory receptors engage with principal vesicular trafficking machineries. CTLA-4 engages with retrograde transport to a substantial extent, more so than the other inhibitory receptors.

**Figure 9.**
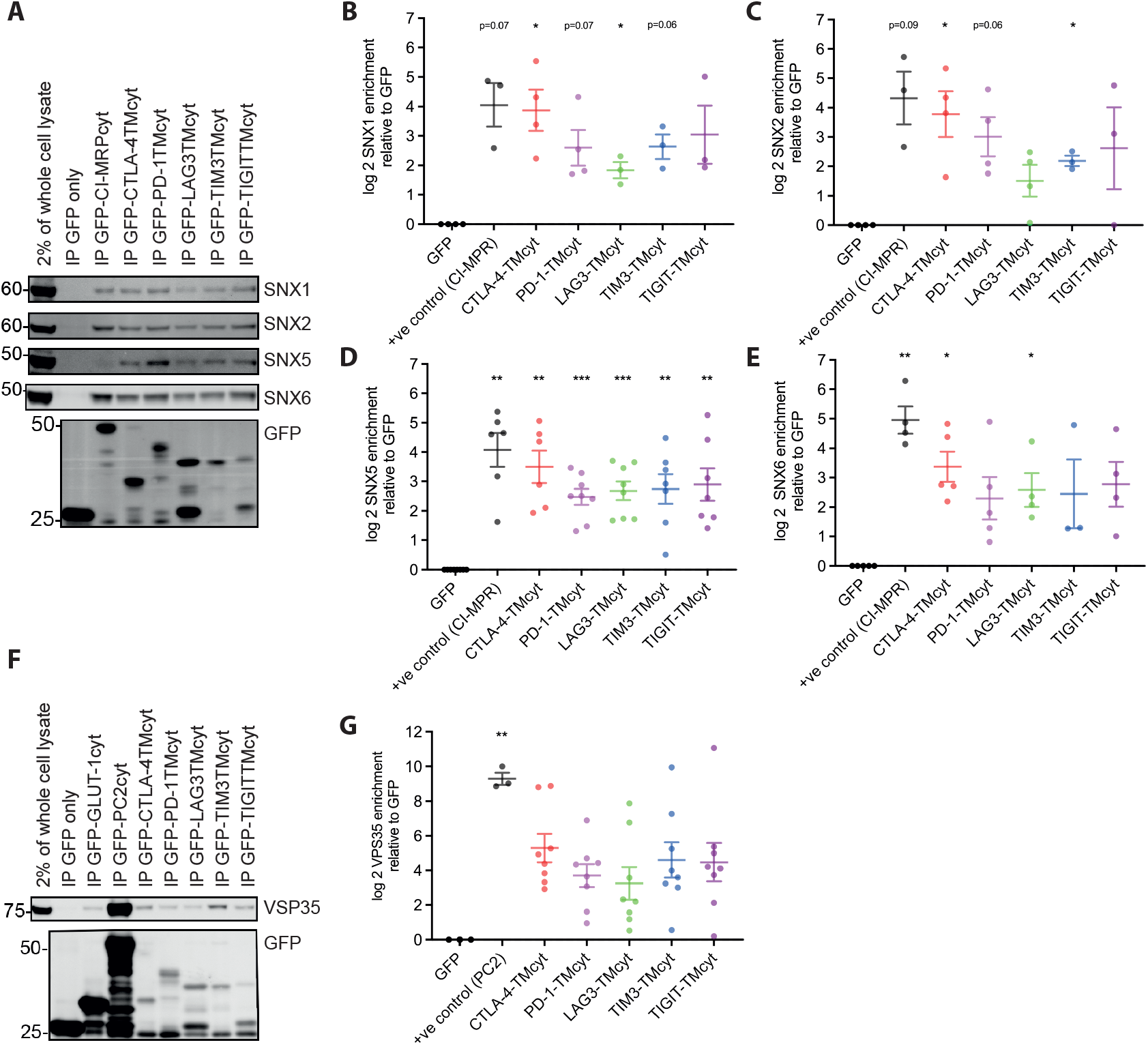
The cytoplasmic and transmembrane domains of inhibitory receptors associate with sorting nexins. **A** Representative Western Blot of GFP pull down experiments from the HEK293T cells in Figure 8F as indicated on top with antibodies against SNX1/2/5/6 and GFP as a control for fusion protein expression as indicated on the right. Full size blots in Figure S7. **B-E** Log2 of enrichment of SNX1 (B), SNX2 (C), SNX5 (D), SNX6 (E) at the cytoplasmic and transmembrane domains of the indicated receptors fused to GFP relative to GFP only. 3 to 8 independent experiments. **F** Representative Western Blot of GFP pull down experiments from the HEK293T cells in Figure 8F as indicated on top with antibodies against VPS35 and GFP as a control for fusion protein expression as indicated on the right. **G** Log2 of enrichment of VPS35 at the cytoplasmic and transmembrane domains of the indicated receptors fused to GFP relative to GFP only. 8 independent experiments. p<0.05, ** p<0.01, *** p<0.001 relative to GFP only; p values calculated using one-way ANOVA

## Discussion

Here we have shown that the majority of the cellular pool of five inhibitory receptors resides in the cell interior (Figure 3) in contrast to four receptors described to be expressed on the cell surface. Comparing the inhibitory receptors to each other in subcellular clustering, the dynamics of plasma membrane insertion, spatiotemporal distribution on the cell surface and molecular protein neighborhood, they were shown to have distinct subcellular distributions. These data are consistent with previous studies showing that different members of receptors families, such as IP_3_ receptors [40], or functionally related groups of receptors, such as ion channels in neurons [41], can show distinct distributions across vesicular structures that adapt to the cellular activation state. Nevertheless, our data on inhibitory receptor distributions in T cells raise important questions.

Which processes limit plasma membrane localization of inhibitory receptors? A likely means to limit plasma membrane expression is constitutive endocytosis. Consistent with this suggestion, the endosomal marker transferrin receptor was highly enriched in the vicinity proteomes of all three inhibitory receptors studied, CTLA-4, LAG3, TIM3 (Figure 6B). Moreover, the endocytic adaptor AP-2 could be pulled down by the transmembrane plus cytoplasmic domains of all five inhibitory receptors (Figure 8I). Association of a non-phosphorylated YVKM motif in the cytoplasmic domain of CTLA-4 is likely to mediate such binding [6,8].

How can distinct vesicular distributions of the five inhibitory receptors be maintained? Conceptually, distinct vesicular distributions can be achieved through receptor-specific recruitment into defined subsets of vesicles or through differential association with the same set of vesicular structures. Given the highly interconnected, dynamic and molecularly complex nature of the vesicular system, these two options may simply be two ends of a continuous spectrum. Even though marker proteins such as EEA-1 or LAMP-1 are commonly used to distinguish different vesicle types, there are extensive contacts between the different vesicle types and of vesicles with microtubules as principal mediators of vesicular trafficking [42–44]. In biochemical fractionation of vesicle types, marker proteins are enriched in single fractions but commonly also appear across a wide range of other fractions [45]. Through consideration of an increasing number of vesicular properties, further vesicle types can be defined in the interconnected and dynamic system, up to 35 in recent imaging-based work [46]. Molecular diversity further blurs the distinction between specific vesicle types and differential distribution across a dynamic continuum of structures. Proteomics identified 1,285 different proteins associated with vesicles in HEK293T cells [47]. An siRNA screen found 132 proteins with 247 interactions between them as strong regulators of endocytosis, 969 proteins with 55,070 interactions as less dominant regulators [48]. Nevertheless, vesicular subtypes with a distinct function can still be defined, such as a subset of endosomes marked by VSP18 that allows the retention of type I interferons for days [49].

It seems unlikely that distinct vesicle subtypes underpin differential localization of the five inhibitory receptors in T cells. Sorting motifs in the inhibitory receptor cytoplasmic domains are overlapping (Figure 8A). Pulldown experiments consistently identified interactions between sorting adaptors and all five inhibitory receptors (Figs. 8, 9), albeit with modest differences in efficiency of association. Colocalization with vesicular markers again showed consistent association of the markers with all four inhibitory receptors studied (Figure 7), again with modest differences in extent. Vesicular proteomes of the three inhibitory receptors studied were distinct, yet with substantial overlap (Figure 6). Consistently differential association with common transport machineries and vesicle types may be sufficient to generate distinguishable subcellular distributions of the inhibitory receptors. Our data also suggest an additional mechanism to generate distinctions in localization. Cargo adaptors can be enriched in specific regions of the plasma membrane [50] and endocytosis can be regulated by cellular adhesion sites [51]. Given the distinct spatial distributions of some inhibitory receptors at the interface between T cells and activating cover slips (Figure 5), an uneven distribution of cargo adaptors and/or differences in the distributions of integrins mediating adhesion [52] may allow sorting of inhibitory receptors into different subsets of endosomes. Different endosome subtypes can then lead to differential downstream sorting [53,54]. Eventually, only a comprehensive characterization of the inhibitory receptor-associated proteomes combined with a functional investigation of the role of the associated proteins in vesicular trafficking will elucidate the mechanisms driving distinct subcellular localization of the five inhibitory receptors. A recent investigation of the TIM3 interactome found 37 high confidence interactors [55]. Thus, a comprehensive elucidation of inhibitory receptor trafficking likely requires substantial effort.

Even without detailed mechanistic understanding of the distributions of the five inhibitory receptors, knowledge of the distinct nature of their distributions is useful. It allows for unique roles of individual inhibitory receptors even in T cells that express all five of them. It thus provides a motivation to carefully evaluate the effects of blocking various combinations of inhibitory receptors on T cell function, for example in checkpoint blockade cancer immunotherapy.

## Supplementary Materials

Figure S1. The extent of subcellular clustering differs between the inhibitory receptors in CD4^+^ T cells

Figure S2. The extent of subcellular clustering differs between the inhibitory receptors in CD8^+^ T cells

Figure S3. PD-1 is largely intracellular in T cells upon lack of TCR engagement

Figure S4. APEX2-mediated biotinylation is specific for vesicles expressing GFP-tagged inhibitory receptors

Figure S5. CTLA-4 is selectively enriched in endosomes late in T cell activation Figure S6. Full size Western blots for Figure 8

Figure S7. Full size Western blots for Figure 9

## Author contributions

Conceptualization, P.J.C. and C.W.; Methodology, S.J.C. and X.R.; Validation, J.L. and C.W.; Formal Analysis, J.L. and C.W.; Investigation, J.L., A.V., B.S., T.R., P.H.C., A.M.P., L.H., P.V. and C.W.; Data Curation, J.L., S.J.C. and C.W.; Writing – Original Draft Preparation, J.L. and C.W.; Writing – Review & Editing, P.J.C. and C.W.; Visualization, J.L. and C.W.; Supervision, A.W.D., D.Y., R.F.M, P.H.C., C.W.; Funding Acquisition, R.F.M., P.J.C. and C.W.

## Funding

Jiahe Lu was supported by a Ph.D. fellowship from the China Scholarship Council. P.J.C. was supported by the Wellcome Trust (104568/Z/14/Z and 220260/Z/20/Z), the Medical Research Council (MR/L007363/1 and MR/P018807/1), the Lister Institute of Preventive Medicine, and the award of a Royal Society Noreen Murray Research Professorship to P.J.C. (RSRP/R1/211004). R.F.M. was supported by the NIH (GM103712).

## Data availability statement

Raw imaging data are available at DRYAD under Raw proteomics data are available at PRIDE under

## Institutional Review Board Statement

Mouse work was approved by the AWERB committee of the University of Bristol under UIN/22/073 on the 19.5.2022.

## Acknowledgements

We acknowledge support from the University of Bristol Flow Cytometry, Wolfson BioImaging and Proteomics core facilities. We thank Maryyam Khabrani for technical support.

## Conflicts of Interest

The authors declare no conflict of interest.

## Methods and Materials

### Mice

CL4 TCR transgenic mice (RRID: IMSR_JAX:005307) [30] were bred and maintained at the University of Bristol under specific pathogen free conditions. CL4 offspring were phenotyped by staining PBMCs with anti-CD8-APC (53-6.7, Biolegend, RRID: AB_312750) and anti-Vβ8.2 FITC (KJ16-133, eBioscience, RRID: AB_465261) antibodies. Homozygous 5C.C7 TCR transgenic mice [28] (RRID:IMSR_EM:13156) were bred and maintained at the University of Bristol under specific pathogen free conditions. BALB/c mice (RRID:MGI:2161072) were purchased from Yuxiu (Shanghai, China). All mice were culled for experimental use between 6 and 12 weeks of age.

### Cells and Media

‘Complete medium’ consisted of RPMI-1640 plus L-glutamine (Gibco) supplemented with 10% FBS (Brazilian source, ThermoFisher), 50µM β-Mercaptoethanol (Gibco), and PenStrep (Gibco) at 100U/mL penicillin and 100µg/mL streptomycin. Renca cells (RRID:CVCL_2174) were maintained in complete medium. CL4 and 5C.C7 T cells were maintained in complete medium supplemented with 50U/mL rh-IL-2 (NIH/NCI BRB preclinical repository)(‘IL-2 medium’). HEK293T (RRID:CVCL_0063) and Phoenix-E cells (RRID:CVCL_H717) were maintained in ‘Phoenix incomplete medium’ consisting of DMEM with 4.5g/L D-glucose, L-glutamine and sodium pyruvate (Gibco) supplemented with 10% FBS (Brazilian source, ThermoFisher), MEM non-essential amino acids (Gibco), and PenStrep (Gibco) at 100U/mL penicillin and 100µg/mL streptomycin. Long-term Phoenix cell stocks were kept in ‘Phoenix complete medium’, which is Phoenix incomplete medium supplemented with 300µg/mL Hygromycin (Invitrogen) and 1µg/mL Diptheria toxin (Sigma).

### Antibodies

Antibodies used are described in the order: antigen, fluorophore if applicable, clone, supplier, dilution, RRID:

For Western Blotting:

SNX1, polyclonal, Abcam, 1:1000, RRID: AB_296721

SNX2, 13, BD Biosciences, 1:1000, RRID: AB_398834

SNX5, polyclonal, Proteintech, 1:1000, RRID: AB_2192708

SNX6, D-5, Santa Cruz Biotechnology, 1:1000, RRID: AB_10842310

SNX27, polyclonal, Proteintech, 1:1000, RRID: AB_10888628

AP-1 γ adaptin, polyclonal, Proteintech, 1:1000, RRID: AB_2058209

AP-2 α adaptin, polyclonal, Proteintech, 1:1000, RRID: AB_2056316

VPS35, polyclonal, Abcam, 1:1000, RRID: AB_10696107

GLUT1, EPR3915, Abcam, 1:1000, RRID:AB_10903230

Polycystin-2, polyclonal, Santa Cruz Biotechnology, 1:1000, RRID: AB_2163376

Anti-mouse IgG, Alexa Fluor 680, Invitrogen, 1:20,000, RID:AB_2633278

Anti-rabbit IgG, Alexa Fluor 800, Invitrogen, 1:20,000, RRID:AB_2633284

For immunostaining and T cell priming:

EAA-1, c45b10, Cell Signaling Technology, 1:200, RRID:AB_2096811

Anti-rabbit IgG, Alexa Fluor 568, Invitrogen, 1:400, RRID:AB_143157

CD3ε, 145-2C11, Biolegend, 5µg/ml, RRID:AB_11149115

CD28, 37.51, BioLegend, 1 µg/ml, RRID:AB_11147170

CD3ε, FITC, 145-2C11, Biolegend, 1:50, RRID:AB_312671

CD4, PerCP/Cy5.5, RM4-5, BioLegend, 1:50, RRID:AB_893326

CD45, APC, 30-F11, BioLegend, 1:50, RRID:AB_312976

CD8α, APC/Fire750, 53-6.7, BioLegend, 1:50, RRID:AB_2572113

CTLA-4, PE, UC10-4B9, BioLegend, 1:50, RRID:AB_313254

PD-1, BV605, 29F.1A12, BioLegend, 1:50, RRID:AB_2562616

LAG3, APC, C9B7W, BioLegend, 1:50, RRID:AB_10639935

TIM3, PE/Cy7, RMT3-23, BioLegend, 1:50, RRID:AB_2571933

TIGIT, BV421, 1G9, BioLegend, 1:50, RRID:AB_2687311

### T cell isolation and stimulation

Procedures used for the isolation and stimulation of TCR transgenic T cells have been extensively detailed elsewhere [29,56–58]. Briefly for CL4 CD8^+^ T cells, red blood cells were removed from dissociated spleens using ACK lysis buffer (Gibco) and splenocytes were plated at a density of 5×10^6^ cells per well, in 1 mL of complete medium, in a 24-well plate. K^d^HA peptide (IYSTVASSL) was added to a final concentration of 1μg/mL and cells were incubated overnight. Primed cultures were stringently washed five times in PBS to remove unbound peptide and then retrovirally transduced (detailed below). Following transduction, cells were re-plated at 4×10^6^ cells per well, in 2 mL IL-2 medium, in a 24-well plate. CTLs were sorted and imaged on day four. For 5C.C7 CD4^+^ T cells, dissociated lymph node cells were plated at a density of 5×10^6^ cells per well, in 1 mL of complete medium, in a 24-well plate. MCC peptide (ANERADLIAYLKQATK) was added to a final concentration of 3µM. T cells were transduced after overnight culture (detailed below). Following transduction, cells were re-plated at 4×10^6^ cells per well, in 2 mL IL-2 medium, in a 24-well plate. T cells were sorted and imaged on day four or five.

For the flow cytometry experiments to quantify endogenous inhibitory receptor cell surface expression, a 24-well cell culture plate was coated with 5μg/mL αCD3 and 1μg/mL αCD28 in phosphate buffered saline (PBS) at 4°C overnight. Red blood cells were removed from dissociated BALB/c spleens using ACK lysis buffer (Gibco) and splenocytes were plated onto a 100mm tissue culture dish for 2 hours at 37°C. Non-adherent cells were taken out and seeded onto the αCD3/αCD28-coated plate at 5×10^6^ cells per well. 48 hours post T cell activation, cells were taken out from the coated plate and stained.

### T cell retroviral transduction

Procedures and plasmids used for retroviral transduction of TCR transgenic T cells have been extensively detailed elsewhere [29,56,57]. Briefly, transfection of Phoenix-E retroviral producer cells with a GFP-sensor expression plasmid was achieved using calcium phosphate precipitation. After transfection cells were cultured for a further 48 hours, then the retrovirus-containing medium was collected and used to transduce T cells. 24h after setting up a T cell culture, T cells were resuspended in 2 mL retrovirus-enriched supernatant from transfected Phoenix-E cells. Protamine sulphate was added to a final concentration of 8μg/mL. Plates were centrifuged for 2 hours at 200xg, at 32°C. Following centrifugation, supernatant was replaced with IL-2 medium.

### Spinning disk confocal imaging

#### Imaging of cell couples

Approximately 72 hours after retroviral transduction T cell cultures were resuspended in ‘imaging buffer’ (10% FBS in PBS with 1mM CaCl_2_ and 0.5mM MgCl_2_) plus DRAQ7. A BD Influx cell sorter (BD Bioscience) was used to isolate GFP-positive T cells [56]. As target cells for CL4 CTL, 1×10^6^ Renca cells were pulsed with K^d^HA peptide at a final concentration of 2µg/mL for 1 hour. These pulsed Renca cells will be referred to as Renca^+HA^ cells. Glass bottomed 384-well optical imaging plates (Brooks life science systems) were used for all imaging experiments. Imaging was done at 37°C using a Perkin Elmer UltraVIEW ERS 6FE confocal system attached to a Leica DM I6000 inverted epifluorescence microscope and a Yokogawa CSU22 spinning disk. A 40x oil-immersion lens (NA=1.25) was used for all imaging experiments, unless otherwise stated. Prior to imaging, 50µL of imaging buffer was added to the well, followed by 5µL of T cells. T cells were allowed to settle for several minutes. Once a monolayer of T cells had formed, 4µL of Renca^+HA^ cells were carefully added to the top of the well. Images were acquired for 15 minutes. Every 20 seconds, a z-stack of 21 GFP images (1µm z-spacing) was acquired, as well as a single, mid-plane differential interference contrast (DIC) reference image. After imaging, GFP-files were saved as individual z-stacks for each time point, and both DIC and GFP files were exported as TIFF files for subsequent analysis.

#### Determination of fluorescence at the cell edge versus interior and of distribution heterogeneity

Fluorescent signal distribution within cells was measured using the ModularImageAnalysis (MIA; version 1.1.1) workflow automation plugin [59] for ImageJ [60,61]. The workflow first detected cells from DIC images using a custom model for the StarDist ImageJ plugin [62]. Detected cells were filtered, with only those falling within an accepted size range retained for further analysis. Each cell was then split in two, based on distance to the edge of the cell, yielding “interior” and “cell edge” cell objects. For analysis, the image slice with the highest mean intensity was extracted from the acquired GFP image stack, passed through a Gaussian filter (sigma = 1 px) and subject to rolling ball background subtraction (radius = 100 px). For each cell interior and edge, the mean and integrated GFP intensity was measured. In addition to this, the intensity distribution within the entire cell was quantified using a greyscale implementation of Ripley’s K-function [63], with measurements taken in the range 1-9 px, at 1 px intervals. Ripley’s K-function compares the distribution of signal at different length scales to that which would be expected by a completely spatially random distribution of the same amount of signal. With this, it is possible to identify length scales at which the signal is either clustering or exhibiting exclusion. Finally, the signal was measured using the Gini coefficient, where the Gini coefficient is a statistical descriptor of the inequality of a population. Coefficient values are given in the range 0 to 1, with these values corresponding to the cases where all pixels have equal intensity and where one pixel has all the signal (and every other has zero signal), respectively.

#### Vesicular reorientation towards interface

Live cell imaging of CL4 T cell MTOC reorientation has been described before [58]. Briefly, DIC files and GFP z-stacks were imported into Metamorph image analysis software (Molecular Devices). For each imaging time point, the GFP z-stack was assembled into a maximum projection. CTLs forming conjugates with Renca^+HA^ target cells were identified, and the time point at which a tight-cell couple forms was assessed using the DIC reference images. Tight cell couple formation was defined as the first time point at which a maximally spread immune synapse forms, or two frames following initial cell contact, whichever occurred first. T cells were divided into three equally wide sections from the cellular interface backwards. If a single cluster of vesicles could be detected, the position of the center of the cluster was recorded as part of one of the three sections.

#### Imaging of individual T cells to estimate association of inhibitory receptors with the cell edge

CL4 CTL retrovirally transduced to express inhibitory receptor GFP fusion proteins were imaged on an Olympus IXplore SpinSR imaging system with a 60x oil immersion lens (NA= 1.50) at 37°C. A z-stack of 65 GFP images (0.25µm z-spacing) was acquired, as well as a single, mid-plane differential interference contrast (DIC) reference image. Association of inhibitory receptor-GFP fluorescence with the cell edge was measured using the ModularImageAnalysis (MIA; version 1.2.7) modular workflow plugin for ImageJ [60,61,64]. The raw image stack was intensity normalised, then cells were enhanced in each slice using a U-Net pixel classification model run via integration with the DeepImageJ plugin (version 2.1.16) [65,66] and a custom model. The resulting probability image was binarized at a probability of 0.5 and both foreground and background regions in the binary image smaller than 200 px^3^ were removed. Cell regions which had become merged in the binary image were then separated using a distance-based watershed transform [67]. Connected components labelling was applied to the binary image stack to create 3D instances of the cells [67] and any detected cells larger than 1800 μm^3^ were discarded as these were assumed to correspond to multiple merged cells which had failed to separate. Any cells in contact with the image edge were also removed from further analysis. Fluorescence measurements were taken on the central slice of each cell, where the membrane is closest to being perpendicular with the imaging plane. First, the segmentation of the central slice of each cell was refined. Multiple radial intensity profiles were extracted, and the refined cell surface taken to be the point at which, when starting from outside the cell, the fluorescence intensity first reached 30%/50% (inhibitory receptors/TCR() of the maximum intensity along that profile. The membrane-to-interior ratio of fluorescence intensity for each refined cell slice was calculated by splitting the cells in two, based on distance from the cell surface. For this, the width of the membrane region was estimated by fitting orthogonal Gaussian profiles around the refined cell surfaces. To estimate membrane width for each cell, the smallest 5/20 (inhibitory receptors/TCR() Gaussian sigma values were averaged and multiplied by 2/3 (inhibitory receptors but TIM3/TIM3, TCR(). The fluorescence intensity within the membrane and interior regions was measured and extrapolated to three dimensions assuming isotropy.

### TIRF imaging

To image vesicular insertion into the plasma membrane, T cells were activated by a flat glass surface coated with anti-CD3ε mAb and soluble ICAM-1. The glass bottom of a 384-well MatriPlate (Brooks Life Sciences) was washed with 50μL 1% acid alcohol (5% (v/v)) 37% HCl in 100% ethanol) at room temperature for 15min and baked at 60°C for up to 30min. 50μL of coating solution (10μg/mL anti-CD3 (Clone 145-2C11, BioLegend) + 2.5μg/mL ICAM1-Fc (Novus Biologicals, #720-IC) was incubated overnight at 4°C. Alternatively, the glass bottom of a 384-well MatriPlate was coated with 1mg/mL biotin-BSA (Sigma-Aldrich), followed by 200µg/mL NeutrAvidin (ThermoFisher) and then 10µg/mL biotin hamster anti-mouse CD3εantibody (clone 145-2C11, BioLegend) and kept in the imaging buffer (1 mM calcium chloride, 500 µM magnesium chloride, 10% (v/v) FBS). No differences in TIRF imaging between the two coating methods were observed, and data were therefore pooled. Live TIRF images were acquired using a Leica AM TIRF microscopy multicolor system attached to a Leica DMI 6000 inverted epifluorescence microscope equipped with 488nm and 561nm laser lines. A Leica HCX PL APO 100x oil immersion lens (NA=1.47) was used. Images were acquired at frame rate every 333ms. For dual color experiments, the laser was switched between frames resulting in image acquisition in each color every 735ms. The onset of T cell activation was determined as the first frame in a series of rapidly spreading TIRF signals.

*To determine the fluorescence distribution across the interface*, a line with width of 20 pixels (4.6µm) was drawn across the widest region of a cell and divided into ten segments. The mean fluorescence intensity of each segment was measured and normalized to segment with the highest fluorescent intensity.

*To determine the frequency of vesicular insertion,* a machine learning-based image analysis algorithm was developed to automatically identify vesicular insertion events at the plasma membrane. The algorithm contained four sequential steps. (i) Blur detection: During time-lapse TIRF imaging, the particle signal might become blurred due to focus drift. To resolve this issue, we developed a support vector machine (SVM) classifier [68] to identify and exclude those blurred frames. As a result, we only retained the sequences of unblurred images in the middle for subsequent analysis. In the classifier, we used a collection of focus measure functions (‘ACMO’, ‘BREN’, ‘CONT’, ‘CURV’, ‘DCTE’, ‘DCTR’, ‘GDER’, ‘GLVA’, ‘GLLV’, ‘GLVN’, ‘GRAE’, ‘GRAT’, ‘GRAS’, ‘HELM’, ‘HISE’, ‘HISR’, ‘LAPE’, ‘LAPM’, ‘LAPV’, ‘LAPD’, ‘SFIL’, ‘SFRQ’, ‘TENG’, ‘TENV’, ‘VOLA’, ‘WAVS’, ‘WAVV’, ‘WAVR’ from ‘https://www.mathworks.com/matlabcentral/fileexchange/27314-focus-measure’) as the feature set. The training data was generated by human annotation of time series images. The parameters in the SVM were optimized using 10-fold cross validation. (ii) Particle detection: The general idea of particle detection is to identify particles using two-sample t-test for local maximum versus local minimum. First, the region of interest (ROI) for particle detection for all time points was determined by 90% of Otsu’s thresholding [69] on the averaged image of all time points (after removing the blurred frames). Then, for each frame, the image was smoothed using wavelet smoothing (Biorthogonal wavelet with 3 and 5 vanishing moments) [70]. After that, the candidate particles were determined as the local maximum within the ROI in the smoothed image; simultaneously, the local minimum points were identified from the smoothed image. Each candidate particle was checked as a valid particle or just noise via two sample t-test. The sample for the candidate point was the pixel values of the five brightest pixels *x_p_* within the 3×3 patch centered by the candidate point. Then, the three closest local minimum points to the candidate point were selected, and the five dimmest points were extracted from each of the 3×3 patches centered on these local minimum points, forming the local minimum sample *x_m_*. After that, the local background standard deviation *σ_l_* with the local maximum sample, and global background standard deviation *σ_l_* with the lower 25% points in the whole frame were calculated. Then, the p-value of *x_p_* − 4*σ_l_* − 3*σ_g_* versus *x_m_* were calculated in the two-sample t-test. If the p-value is below a predetermined threshold (0.001 in this case), the point is considered a valid particle; otherwise, it is classified as a noise point. This process is repeated for all candidate points and all frames. (iii) Point tracking. After point detection, we used the simpletracker library (https://www.mathworks.com/matlabcentral/fileexchange/34040-simpletracker) to track all the particles over time. We used the nearest neighbor algorithm in the linkage with linkage threshold of 2√2, and a gap threadhold of 3 frames. (iv) Event classification. Tracked particles were classified into two categories. ‘Vesicle insertion’ events were characterized by rapid lateral dispersion of the fluorescence signal of a newly detected particle (Figure 4B). ‘Microcluster’ events were stable. We don’t know what these events represent, possibly receptor microclusters. They were excluded from the subsequent counting of vesicle insertion events. After particle detection and tracking, we determined the event type with machine learning classification and prediction. The feature is the log10 of the p-value of each particle over the local background via two sample t-test. The training data was generated by human annotation of time series images. The SVM classifier was trained on the annotated data with 10-fold cross validation. The events are predicted for each particle at each time point. The output file of this analysis was a list of events with onset time, duration, coordinates within the entire imaging field and classification.

*For colocalization analysis in dual color TIRF experiments,* colocalization over time was measured using a workflow compiled in the ModularImageAnalysis (MIA, version 0.21.13) workflow automation plugin [71] for Image J [60,61]. Cells in the two fluorescence channels were segmented starting with a 2D median filter (radius = 2 px) followed by rolling ball background subtraction (radius = 50 px). Filtered images were then binarized using local thresholding with the Sauvola algorithm (radius = 50 px) [72] and any holes or isolated fragments in the binarized images smaller than 25 μm^3^ were removed. Of the remaining regions in the binary images, only those with 2D circularity greater than 0.2 were retained. The binary images from the two input channels were merged and subject to an additional 2D median filter (radius = 2 px). Any remaining binary regions which had become merged were split apart using a distance-based watershed transform [67]. Individual cell objects were identified from the binarized images using connected components labelling [67] and any cells falling outside a 50 μm^3^ to 500 μm^3^ accepted size range were discarded. Cells were tracked through time [73], and any tracks detected in an insufficient number of frames were also discarded. Pearson’s Correlation Coefficient was measured within each cell object for the two, raw fluorescence image channels.

### APEX proteomics and electron microscopy

*APEX proteomics:* 2×10^6^ CL4 T cells transduced and FACS sorted to express CTLA-4-GFP, LAG3-GFP or TIM3-GFP were incubated with 500μM biotin-phenol (Iris Biotech) in IL-2 medium at 37°C for 30 minutes. Cells were collected and incubated with 1mM H_2_O_2_ (Sigma-Aldrich) for exactly 30s and spun for 30s at 311g. Cells were immediately washed four times with 200μL quenching buffer (10mM Sodium Azide, 10mM Sodium Ascorbate, 5mM Trolox in 0.1M PBS), once with 0.1M PBS and left on ice. Cell pellets were then lysed at 4°C with 80μL lysis buffer (50mM Tris, 0.5% (v/v) NP40, 1% (v/v) protease + phosphatase inhibitor cocktail) for 20min. To the 20μL lysis buffer prewashed streptavidin beads (GE Healthcare) were added per sample and rotated for 2h. The beads were washed twice with wash buffer (50 Mm Tris, 0.25% NP40, 1% Protease + Phosphatase inhibitor cocktail) and then wash buffer without NP40 then processed for proteomic analysis. Biotinylated proteins were digested into peptides and labelled with TMT 6-plex tandem mass tags. Using liquid chromatography–tandem mass spectrometry, peptide sequence information was obtained by an MS1 scan, and then the reporter group of the TMT 6-plex labels was cleaved to detect their specific m/z by MS/MS whose intensity represents the relative amount of the peptide from the corresponding samples [74]. Mass spectrometry data were analyzed with seaMass, a protein-level quantification and differential expression analysis method [75]. seaMass generates protein-level quantifications for each sample through Bayesian mixed-effects modelling [76] accounting for variation due to digestion and sample-level variation and automatically infers normalisation effects across and within iTraq/TMT plexes. To account for the correlation between paired positive and negative samples an additional random effect was included in the experimental design. Differential expression estimates of the log-2-fold change of protein expression between each of the CTLA4, TIM3 and LAG3 conditions versus the negative control condition were generated. Seamass’s inferred per-sample protein-level quantifications were then used to generate t-sne plots [77] which shows the similarity of the protein-level quantifications between samples.

*APEX electron microscopy:* 3×10^6^ Renca cells were pulsed with 2μg/mL K^d^HA peptide for 1h. At least 1×10^6^ APEX2-CTLA-4/LAG-3/TIM-3-GFP overexpressing CL4 T cells were sorted by FACS. Both cell types were suspended in 100μL 10% FBS/PBS/1mM CaCl_2_/0.5mM MgCl_2_, mixed and spun 30s at 311 g. The supernatant was removed, and the cell pellet was incubated at 37°C for 5min. The cell pellet was fixed in 2.5% glutaraldehyde/3mM CaCl_2_/0.1M sodium cacodylate, pH7.4 for 5min at room temperature, then 1h on ice. The cell pellet was washed with 0.1M sodium cacodylate, pH7.4 (wash buffer), blocked in 30mM glycine in wash buffer, then washed again in wash buffer, each step for 10 min. The wash buffer was replaced with a DAB + 0.03% H_2_O_2_ solution for 5min. Cells were washed 10min in wash buffer, followed by 30min incubation with 1% osmium tetroxide. The cell pellet was washed with distilled water, fixed in 3% uranyl acetate for 20min, washed with distilled water, and dehydrated through 70%, 80%, 90% 96% and 3 times 100% ethanol, 10mins each. The pellet was infiltrated in 100% EPON resin rotating overnight at room temperature, then embedded in fresh 100% EPON resin, polymerized in 12-24 hours at 65°C. Samples were cut into 20μm sections and imaged by transmission electron microscopy.

### Immunostaining

#### Indirect immunostaining for EAA1

20,000 Renca cells in 50µL complete medium were seeded in a 384-well plate (Corning) and incubated overnight at 37°C. Renca cells were incubated with 1µg/µL of HA peptide for 1h. 100,000 CL4 T cells transduced with GFP-tagged co-inhibitory receptors (LAG-3, TIM-3, CTLA-4 or PD-1) were added and left for 10min at 37°C. Cells were fixed in 50µL 4% PFA for 20min at 4°C. Each well was washed with 50µL of PBS. The reaction was quenched with 50µL of 50mM NH_4_Cl for 10 minutes at 4°C, and wells were washed three times with 50µL PBS. Cells were permeabilized using 50µl of 0.02% Triton X-100 in PBS for 20min at 4 °C. Cells were washed with 50µl of PBS and incubated with 50µl of blocking solution (1% BSA in PBS) for 30min at room temperature.

Cells were stained with 50µl anti-EEA1 mAb (Cell Signaling Technology) at a dilution of 1:200 in 1% BSA in PBS with Fc Block at the same dilution overnight at 4 °C. Cells were washed three times with 50µL of PBS. The wells were incubated with 50µl of secondary antibody (Anti-Rabbit IgG, Alexa Fluor 568) diluted 1:400 in 1% BSA in PBS with Fc Block at 1:200 for 1 h at room temperature. Wells were washed three times and left in 50µL of PBS at room temperature. Nuclei were stained with Hoechst 33258 (Invitrogen) diluted 1:5000 in PBS for 10min at room temperature.

#### Lysotracker/Mitotracker staining

20,000 Renca cells in 50µL complete medium were seeded in a 384-well plate (Corning) and incubated overnight at 37°C. Renca cells were incubated with 1µg/µL of HA peptide for 1h. 300,000 CL4 T cells transduced with GFP-tagged co-inhibitory receptors (LAG-3, TIM-3, CTLA-4 or PD-1) were suspended in IL-2 medium at 37°C containing either 65nM LysoTracker® Deep Red (ThermoFisher Scientific) or 25nM MitoTracker™ Red CMXRos for 1h or 30min, respectively. Following incubation, cells were resuspended in 37°C IL-2 medium. 100,000 CL4 T cells were added and left for 10min at 37°C. Cells were fixed and stained with Hoechst 33258 as described in the EEA1 staining protocol.

#### Image acquisition

All images were collected with a Leica SP8 AOBS confocal laser scanning microscope attached to a Leica DMi8 inverted epifluorescence microscope. A HC PL APO CS2 40x oil immersion objective (NA=1.3) was used. GFP constructs were excited with 488nm line of the white light laser, Hoechst-stained nuclei with the 406nm line of the diode laser and LysoTracker/EEA1/MitoTracker with the 561nm line of the white light laser. The pinhole was set to 1 airy-unit (AU). A zoom factor of 6 was used to acquire all the images.

#### Colocalization analysis

All images were analyzed using the Fiji-ImageJ [60,61] software to determine the Pearson’s correlation coefficient (PCC) for overlap between EEA1/LysoTracker/Mitotracker and GFP-tagged co-inhibitory receptors.

Sets of images without Hoechst stain were analyzed using an image analysis workflow created using MIA plugin (v0.18.2) for the Fiji-ImageJ software: To determine the best focus slice from the Z-stack, the GFP channel was filtered with a Difference of Gaussian (DoG) filter (radius=6px) to enhance the bright band around the edge of the cell. The Fiji Ridge Detection plugin was used to detect ridge-like objects [58,78]. The best focus slice was identified as the slice with the most “ridge” features – longest summed ridge length. To identify membranes in the images, images were converted to RGB format. Pixel classification was applied using the WEKA Trainable Segmentation plugin to enhance membrane structures [79]. A 2D median filter (radius=2px) was applied to the membrane probability image, based on the GFP signal. The image was binarized using a global Otsu threshold. Foreground and background regions smaller that 300px^3^ in volume were removed [67]. An 2D erode binary operation (radius=2px) was applied to shrink objects.

Membranes were detected as foreground-labelled pixels [67]. One analyzable object from each image was detected (includes all cells qualifying for co-localization analysis). The PCC was calculated for detected membrane regions as detailed below.

Sets of images with Hoechst stain were analyzed using an image analysis workflow created using MIA plugin (v0.18.15) [80] for Fiji-ImageJ software [60,61]: To detect nuclei, a 3D median (radius = 2px) was used to smooth the Hoechst 33258 image whilst retaining sharp edges on the nuclei. A 2D sliding paraboloid background subtraction (radius=30px) was performed on all z slices. The image was binarized using the global Huang threshold [81], followed by 2D hole filling and 2D median filtering (radius=5px). Nuclei were initially identified as contiguous regions of foreground-labelled pixels using connected components labelling [67]. Any objects detected with volumes less than 20µm^3^ are excluded. Initial objects were updated to correspond to all pixels enclosed with an alpha shape (calculated using the MATLAB implementation) [82]. Adjacent nuclei objects that have become merged during binarization were separated using a distance-based watershed transformation [67]. To calculate the PCC for detected single cell membrane regions, nuclei were removed from the foreground of the binarized cell image, leaving just the cytoplasm. The binarized cell image was sub-divided, such that all points of a contiguous region were closest to the same nucleus. Using connected components labelling, cytoplasm objects were detected in the sub-divided, binarized images [67]. Red and green-channel intensities were measured for all points of the detected cytoplasm objects, and any objects with mean green-channel intensity lower than a user-defined threshold were excluded from further analysis. PCC was calculated for each object using the red and green channels.

#### Immunostaining to determine fraction of an inhibitory receptor on the cell surface

3×10^6^ day 3 BALB/c T cells were stained for surface receptors in 1% BSA/PBS for 30min at room temperature. The surface-stained cells were washed twice with PBS. 1/3 of the cells was directly analyzed by flow cytometry; 2/3 of the cells were fixed with 750μL fixation buffer (BioLegend, #420801) for 30min, washed twice with 1x permeabilization buffer (10% (v/v) in ddH_2_O, BioLegend, #421002), stained again in 1x permeabilization buffer for 30min at room temperature and analyzed by flow cytometry.

### Immunoprecipitation

Based on the pEGFP.C1 vector (Clontech) plasmids for the CMV promoter-driven expression of fusion proteins of GFP with either only the inhibitory receptor cytoplasmic domain or the joint transmembrane plus cytoplasmic domain were generated. HEK293T cells were grown to 90% confluence in 15cm culture dishes and transfected with 15μg DNA of GFP-inhibitory receptor cytoplasmic domain or transmembrane plus cytoplasmic domain constructs using 1μL of a 10mM polyethylenimine stock solution in 5mL Opti-MEM I reduced Serum Medium (Life technologies, #31985). GFP-trap beads (Chromotek, GTA20) were aliquoted to 1.5μL per sample and equilibrated by three washes in IP lysis buffer (50mM Tris/HCl, 0.5% NP40, complete protease inhibitor cocktail (ThermoFisher, #A32955), pH=7.4). HEK2393T cells were washed twice with ice-cold PBS and lysed by prechilled IP lysis buffer on ice and centrifuged at 13,000 g for 10 minutes at 4°C. 2% of the cell lysate supernatant was kept as input sample and the rest of the supernatant was incubated with the equilibrated GFP-trap beads for 1h at 4°C on a rotator. GFP-trap beads were washed three times with IP lysis buffer and once with IP lysis buffer without NP-40. Proteins were eluted from the beads and denatured in 2x sample buffer (4% SDS in lysis buffer) at 95°C for 10 minutes. Proteins were detected by Western blotting using a LI-COR Odyssey scanner.

## Notes

### Competing Interest Statement

The authors have declared no competing interest.

### Summary of Updates

correction of an author middle name

